# Estimating densities of larval Salmonflies (*Pteronarcys californica*) through multiple pass removal of post-emergent exuvia in Colorado rivers

**DOI:** 10.1101/2019.12.13.875195

**Authors:** Brian D. Heinold, Dan A. Kowalski, R. Barry Nehring

**Author notes:** Corresponding author, (DK).

## Abstract

Traditional methods of collecting and sorting benthic macroinvertebrate samples are useful for stream biomonitoring and ecological studies; however, these methods are time consuming, expensive, and require taxonomic expertise. Estimating larval densities through collection of post-emergent exuvia can be a practical and time efficient alternative. We evaluated the use of multiple pass depletion techniques of the post-emergent exuvia of *Pteronarcys californica* to estimate larval densities at ten sites in three Colorado rivers. Exuvia density was highly correlated with both final-instar larval density (R^2^ = 0.90) and total larval density (R^2^ = 0.88) and the multiple pass removal technique performed well. Exuvia surveys found *P. californica* at three low density sites where benthic sampling failed to detect it. At moderate and high density sites the exuvia surveys always produced lower density estimates than benthic surveys. Multiple pass depletion estimates of exuvia proved to be an accurate and efficient technique at estimating larval densities and provided an effective alternative for traditional benthic sampling when objectives are monitoring *P. californica* and detecting populations, especially at low density sites.

## Introduction

Evaluating the condition of freshwater ecosystems through benthic macroinvertebrate communities is a common approach for stream health assessment and biomonitoring [1-3]. These methods characterize and compare aquatic invertebrate communities among sites using regionally developed standards. Benthic studies, while useful, are labor and time intensive, expensive, sensitive to sampling techniques, and require taxonomic expertise. The costs can be justified by the valuable data used by government agencies, researchers, and water managers to maintain and monitor water quality and understand function of river ecosystems. But, if sampling objectives are more specific and budgets are limited, whole community benthic sampling may not be necessary or the most appropriate technique.

One ecologically important aquatic invertebrate commonly used as a bioindicator is the Giant Salmonfly (*Pteronarcys californica* Newport). It is useful for biomonitoring because of its sensitivity to habitat alteration, widespread distribution in western North America [4, 5], multi-year larval life stage, large body size, easy identification, low larval dispersal, and well defined larval habitat preferences [6-9]. *Pteronarcys californica* is among the largest and longest lived stonefly in western North America [10-12]. In Colorado, larvae typically inhabit unpolluted, medium to large, permanent streams with unconsolidated cobble and large gravel substrates between 1,500 and 2,500 m in elevation [13, 14]. Adults emerge from late May to early July and recruitment begins in April after a 9-10 month egg diapause [15] followed by a three to four year aquatic larval stage [16, 17]. Mature larvae (larvae expected to hatch that year) migrate toward the stream bank to stage a highly synchronous adult emergence. Salmonflies typically emerge at night crawling out of the water onto riparian substrates to become winged terrestrial adults where they leave post-emergent exuvia (hereafter, exuvia).

*Pteronarcys californica* plays an important ecological role, both in biomass and abundance, in stream and riparian food webs. As shredders, larvae process coarse organic matter like vascular plants and algae [9, 18] making the nutrients available to other feeding groups as detritus or body biomass [19]. Salmonflies can comprise a large portion of the benthic biomass because of their large body size and high densities in suitable habitat [20, 21], making them an important component of stream food webs for crayfish, other invertebrates, and trout [22, 21]. Terrestrial adults are part of a critical link for aquatic-riparian nutrient and energy exchange [23] as prey for frogs, birds, bats, and spiders [15, 24]. Despite its ecological importance, range-wide declines of *P. californica* have been documented in the Logan and Provo Rivers in Utah [25, 26], several rivers in Montana [27], and in the Gunnison and Colorado Rivers in Colorado [4, 28] mostly due to effects of dams like decreased water quantity and quality, siltation, and pollution.

Density of benthic macroinvertebrates is traditionally estimated by systematically collecting samples from a fixed area of the stream bed. Alternative methods have been recently developed to indirectly survey communities by identifying and enumerating exuvia. These methods can reduce time and labor of traditional techniques while providing reliable population density estimates, community structure, and life history information. Ruse [29] deduced chironomid communities from larval and pupal exuvia and Foster and Soluk [30] estimated densities of the endangered Hine’s emerald dragonfly (*Somatochlora hineana*) more accurately by sampling larval exuvia than by collecting adults. Raebel et al. [31] stated the importance of exuvia collections to avoid bias in adult Odonata surveys. DuBois [32] enhanced these studies by using a depletion population estimator to approximate exuvia densities and detection probabilities of Anisoptera. Richards et al. [33] correlated *P. californica* exuvia densities and live (wet) larval body weights with substrate embeddedness to demonstrate differences in life history, distribution, and abundance above and below a main stem impoundment. Their work provided a foundation in the development of our novel technique to estimate larval densities through multiple pass removal sampling of exuvia.

Multiple pass removal sampling is a commonly used technique in wildlife and fisheries to estimate population size of closed populations. Assumptions of this models used to analyze these data are closure (no deaths, births, emigration, or immigration) and constant capture probability [34, 35] that must be met to avoid bias [36, 37]. If more than two depletion events are completed then assumptions about capture probabilities can be relaxed and capture rates for different passes can be estimated. If populations can be considered geographically and demographically closed (due to isolation or short sampling time period) then population estimation can be accomplished rather simply if good unbiased estimates of detection probability are possible.

The objective of this study was to couple traditional benthic invertebrate sampling with multiple pass removal techniques to evaluate if closed population estimation models can be used to estimate the density *P. californica* larvae. We tested this by correlating densities of systematically collected exuvia from the riparian area with densities of larvae from benthic samples. Another goal was to provide a safer and more efficient method for estimating single species densities.

## Methods

### Study area

Benthic and exuvia sampling was conducted at ten sites on three rivers in Colorado. Eight sites were sampled on the Colorado River and one on the Fraser River both in north central Colorado, and one site on the Gunnison River in southwest Colorado (Fig 1). Distance between the lowest Colorado River site and the Fraser River sites is 74 km.

**Fig 1.**
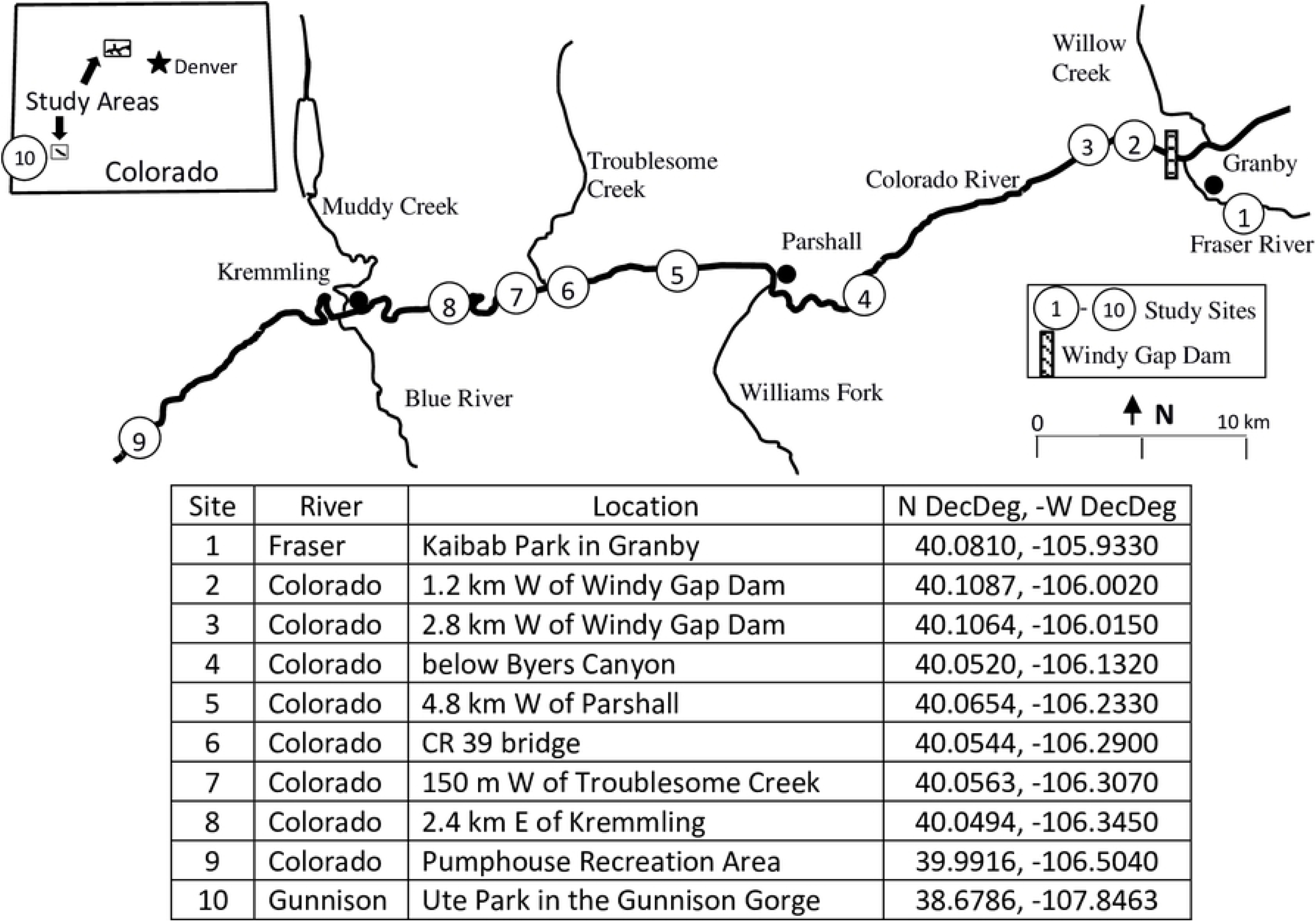
*Pteronarcys californica* benthic and exuvial collection sites in 2010 from the Colorado and Fraser Rivers. Gunnison River site shown only on inset map.

### Benthic sampling

Three benthic subsamples were taken at each site between 15 - 18 April 2010 from the Colorado and Fraser Rivers and 10 May 2010 from the Gunnison River, approximately 1 month prior to the typical adult emergence times of *P. californica*. All sites were located in riffle areas dominated by cobble substrates interspersed with gravel except for sites 7 and 8 which were dominated by sand and gravel. A modified Surber sampler with a 0.25 m^2^ sampling frame (55.0 cm x 45.5 cm) and 150 μm mesh net was used. Cobbles larger than 10 cm in diameter were individually scrubbed with a brush, invertebrates washed into the net, and then the cobbles removed from the sampling frame. Remaining substrate within the frame was disturbed to a depth of 10 cm to dislodge invertebrates into the net. Contents were preserved with 80% ethanol in 2 L plastic jars.

In the lab, all *P. californica* larvae were sorted, sexed [21, 38], and measured for total length (TL) from the anterior tip of head to the posterior tip of the epiproct to the nearest millimeter under a dissecting microscope with a calibrated ocular micrometer. Length frequency histograms for male and female larvae were constructed based on TL to separate annual year classes. Densities of mature larvae and densities of all larvae were calculated and used in separate analyses for correlation with exuvia densities. Mature larval cut off lengths were distinct from the younger year class providing reliable data for analysis despite problems with TL measurements. Separating cohorts and year classes of merovoltine species has proven difficult because of varying growth rates [16] and contraction or expansion of abdomens in preserved insect specimens can further confound this task. Our colleagues [39] used head capsule width and combined head and thorax lengths to produce “body size” or “body area” to assign cohorts within a stream.

### Exuvia sampling

Sampling began with the onset of *P. californica* adult emergence on the Colorado River at site nine on 2 June 2010 and proceeded upstream to end at site one on the Fraser River on 21 June 2010; sampling at site 10 on the Gunnison River lasted from 16-23 June 2010. Each site was sampled beginning on the day when the first exuvia was found or winged adults were observed and continued daily until exuvia were no longer found. Data collection was performed by searching for exuvia within 10 m of the bank along two 30.5 m transects on one side of the river directly adjacent to benthic sampling sites. Collections at a site were accomplished by 2-4 people in a matter of 2-6 hours completing 2-4 passes with identical effort and personnel. Specimens were taken only when attached to dry riparian and emergent substrates; none were taken from the water to avoid counting ones that possibly drifted into the site. Exuvia were enumerated on hand held counters, stored in sealable bags, and removed from the search area. Amount of time searching varied by site depending on the number of exuvia and complexity of riparian search area.

### Data analysis

Area of benthic habitat was estimated by multiplying the sampling section length (always 30.5 m) by the average wetted channel width derived from 10 evenly spaced cross-channel transects. To evaluate the assumptions of the removal model and appropriateness of this sampling technique, three and four pass removal data were compared to two pass data for twenty of the sampling events. Three and four pass data were analyzed with the Huggins Closed Capture model in Program Mark [40, 41] and two pass data were analyzed with the simpler two pass removal model [34]. In Mark, models were built that varied capture probability by pass, allowing a different capture probability for the first pass and the second pass and the third pass or third and fourth passes. Declining capture probability with subsequent passes is a common source of bias of removal models in fisheries data [36, 37] and comparing the population estimates and capture probabilities allowed us to evaluate the assumption of constant capture probability of the simpler two pass model. The assumptions of demographic and geographic closure were expected to be met due to immobility of exuvia and the emergence occurring at night. To evaluate if exuvia densities accurately estimated larval densities, we used simple linear regression in R [42]. Exuvia densities were the dependent variable and densities of mature larvae and all age class larvae were both used in separate analyses as the independent variable.

## Results

Adult emergence of *P. californica* lasted between 2-8 days at each site and proceeded upstream approximately 4 km per day. Early in the emergence, male exuvia were dominant and sex ratios were more even toward the end of the emergence. Approximately 97% of exuvia (n=21,526) were collected within 2 m of the bank. A total of 592 larvae were collected. Larvae from the Colorado and Fraser Rivers separated into four year classes; mature female larvae were ≥39 mm TL (mean 46.5, SE 0.51) and males ≥ 35 mm TL (mean 39.2, SE 0.34) (Table 1). Larvae from the Gunnison River separated into three year classes; mature female larvae ≥41 mm TL (mean 49.1, SE 0.51) and mature males ≥ 37 mm TL (mean 41.9, SE 0.46) (Table 2). Mature females were significantly larger than mature males within each river (p=0.0000 for each).

**Table 1.** Year class lengths and frequency distribution of *Pteronarcys californica* larvae collected 30 April- 1 May 2010 from the Colorado and Fraser Rivers. Lengths in mm from anterior tip if head to posterior tip of epiproct.

| Year Class | Male larvae<br>(n=149) | Female larvae<br>(n=123) |
| --- | --- | --- |
| 1 | $\leq 15$ mm (33) | $\leq 17$ mm (41) |
| 2 | 16-25 (34) | 18-25 (25) |
| 3 | 26-34 (31) | 26-38 (29) |
| 4 | $\geq 35$ (51) | $\geq 39$ (28) |

**Table 2.** Year class lengths and frequency distribution of *Pteronarcys californica* larvae collected 14 April 2010 from the Gunnison River. Lengths in mm from anterior tip if head to posterior tip of epiproct.

| Year Class | Male larvae<br>(n=191) | Female larvae<br>(n=129) |
| --- | --- | --- |
| 1 | $\leq 23$ mm (162) | $\leq 23$ mm (95) |
| 2 | 24-36 (14) | 24-40 (12) |
| 3 | $\geq 37$ (15) | $\geq 41$ (22) |

Exuvia densities were highly correlated with both mature larval densities (R^2^ = 0.90) and total larval densities (R^2^ = 0.88). Exuvia densities averaged 2.6/m^2^ and ranged from 0.002/m^2^ to 11.443/m^2^ (Table 3). Total larval density averaged 80.0/m^2^ and ranged from 0 to 437.3/m^2^. Density of mature larvae averaged 16.1/m^2^ and varied from 0 to 101.3/m^2^. The correlation between mature larvae and exuvia densities was high but the relationship was not 1:1. Larval estimates were generally higher than exuvia estimates except at sites 2, 3, and 7 where exuvia were found but no larvae. To predict the density of mature larvae, the linear equation was: larval density = 7.358*(exuvia density) - 2.854.

**Table 3.** Densities in m^2^ of *Pteronarcys californica* pre-emergent larvae, all larvae, exuvia, and population estimates from exuvia collected from April-June 2010 from the Colorado, Fraser, and Gunnison Rivers.

| Site | Densities m<sup>2</sup> |  |  |  |
| --- | --- | --- | --- | --- |
|  | All larvae | Pre-emergent larvae | Exuvia | Exuvia population estimate |
| 1 | 8.00 | 1.33 | 0.854 | 0.872 |
| 2 | 0.00 | 0.00 | 0.064 | 0.065 |
| 3 | 0.00 | 0.00 | 0.077 | 0.079 |
| 4 | 4.00 | 4.00 | 2.391 | 2.477 |
| 5 | 4.00 | 1.33 | 0.686 | 0.697 |
| 6 | 44.00 | 4.00 | 0.306 | 0.315 |
| 7 | 0.00 | 0.00 | 0.002 | 0.002 |
| 8 | 5.33 | 0.00 | 0.007 | 0.007 |
| 9 | 297.33 | 101.33 | 11.001 | 11.443 |
| 10 | 437.33 | 49.33 | 9.392 | 9.849 |

Exuvia detected populations at all 10 sites whereas larvae were found at only seven of the 10 sites (Table 3). Capture probabilities of exuvia ranged from 0.45 to 0.89 (average 0.72). Simple two pass population models were sufficient to produce unbiased population estimates. Capture probabilities and population estimates were very similar for both the Huggins closed capture model for three and four pass estimates and the Zippin two pass estimates (Fig 2). The two pass depletion technique worked best at sites with moderate exuvia densities and there was some variation in capture probability at very low densities (<80 exuvia per 30.5 m) and very high densities (> 6,000 exuvia per 30.5 m) indicating that the assumption of equal capture probabilities for all passes is violated with the simple two pass model. However, that bias was small and population estimates of the two models were very close (Fig 2).

**Fig 2.**
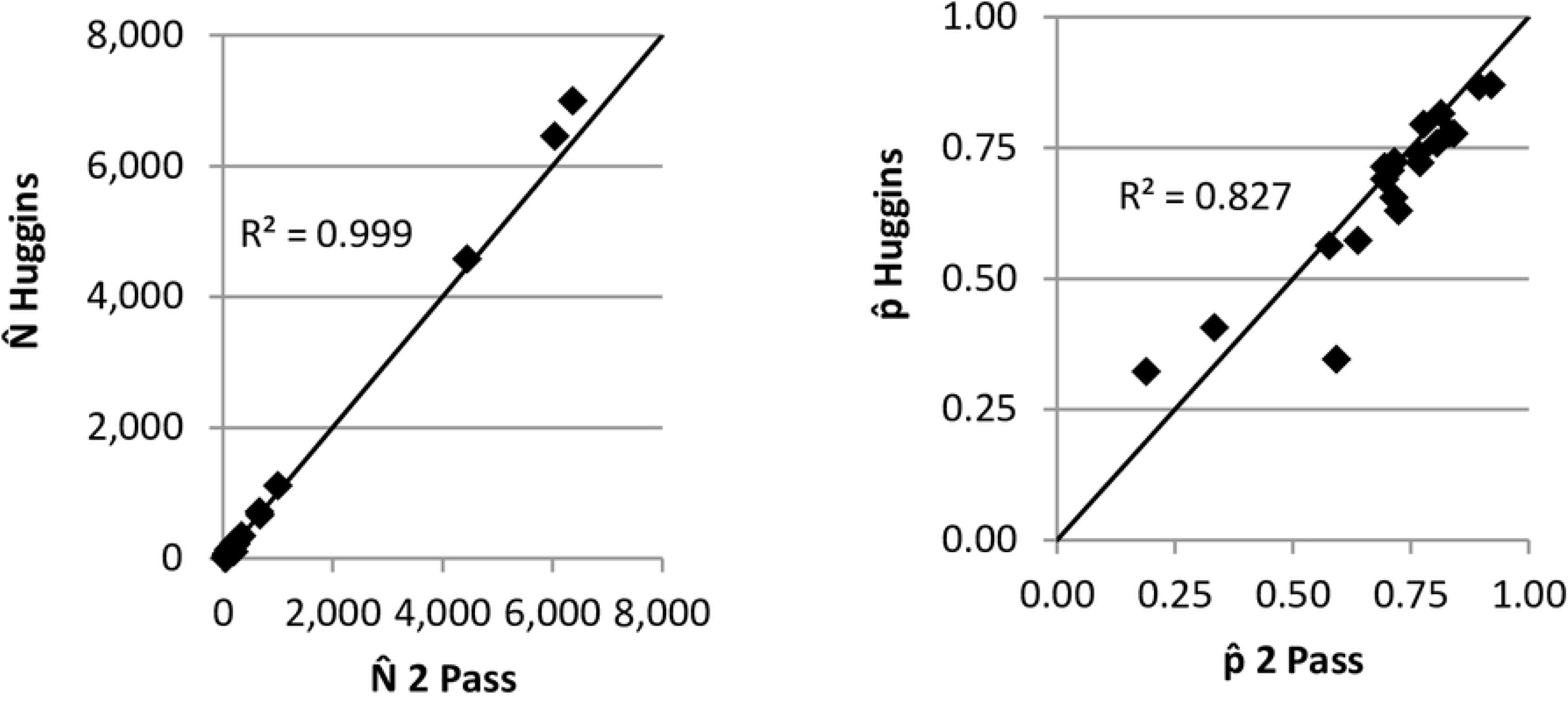
Population estimates 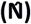 and capture probabilities 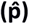 of a three pass Huggins Closed Capture model in Program Mark (with time effects that varied capture probabilities) and a simple two pass removal model of Zippin 1956.

## Discussion

Multiple pass removal estimates of *P. californica* exuvia effectively predicted densities of mature larvae. Assumptions of the multiple pass depletion models appeared to be met and capture probability varied minimally among passes. The two-pass depletion technique performed well due to immobility of exuvia, high capture probability, and no size selective gear [36, 37, 43], suggesting two sampling passes can be adequate if three or four passes are cost prohibitive as with Odonata exuvia [32].

Correlation between densities of exuvia and mature larvae were high but not 1:1. Exuvia underestimated larvae at high density sites but at low density sites exuvia overestimated larvae. The underestimation of larval densities is likely attributed to the behavior of mature larvae congregating near the river bank prior to emergence in the shallow, wadeable water where benthic sampling must occur, creating an artifact of unnaturally high densities. Other factors that may contribute to an underestimate are imperfect detection probability of exuvia and dispersal of larvae out of the sampling area or predation in the 1-2 month time period between benthic sampling and emergence. Overestimates were likely due to the high capture probability of exuvia in addition to the difficulty of collecting larvae that are rare at a site using Hess or Surber samplers [44]. Therefore, exuvia sampling may more accurately estimate, not necessarily overestimate, larval densities than benthic sampling at low density sites.

Detection rates of populations through exuvia sampling were higher than for benthic sampling. This is likely because the large amount of available benthic habitat was essentially reduced to a much smaller, well defined and more easily accessible riparian sampling area. Riparian sampling area among sites averaged 61 m^2^ (30.5 m long x 2 m wide) compared to 742 m^2^ (400-1500 m^2^) of unevenly distributed benthic habitat, much of which may not be accessible by wading due to excessive depth or water velocities ≥ 2 m/s.

Presence of exuvia or adults is the only evidence of successful life cycle completion. Varying densities can indicate habitat quality and help identify reference sites and priority areas for river conservation, restoration, and monitoring of *P. californica*. In regions where *P. californica* does not occur, this technique may be useful for other easily recognizable stoneflies like *Pteronarcella badia, Claassenia sabulosa, Hesperoperla pacifica*, or mayflies like *Timpanoga hecuba*. This technique eliminated the need for benthic sample collection, preservation, and subsequent expense of processing in the laboratory. It also provided accurate and less biased density estimates of *P. californica* larvae than those derived from benthic samples.

Benthic sampling of aquatic invertebrates is a useful and productive biomonitoring technique but the overall process to acquire data can be labor and cost intensive. In addition, it can be difficult to find target species that are rare at a site with benthic sampling [44]. Using multiple pass removal sampling of the recently shed exuvia can be an effective and efficient way to estimate densities of *P. californica* and may be superior to traditional benthic sampling at detecting the species at very low densities.

## Acknowledgements

We would like to thank Justin Pomeranz, Jon Ewert, technicians, and volunteers for tireless field work counting exuvia. Dr. Boris Kondratieff kindly provided valuable lab space and equipment.

## References

1. Goodnight CJ. The use of aquatic macroinvertebrates as indicators of stream pollution. Transactions of the American Microscopical Society. 1973; 92: 1–13.

2. Rosenberg DM, Resh VH. Introduction to freshwater biomonitoring and benthic macroinvertebrates. Chapman and Hall, New York. 1993.

3. Barbour MT, Gerritsen J, Snyder BD, Stribling JB. Rapid bioassessment protocols for use in streams and wadeable rivers: periphyton, benthic macroinvertebrates and fish. 2nd edition. U.S. Environmental Protection Agency; Office of Water; Washington, D.C. 1999

4. Elder JA, Gaufin AR. Notes on the occurrence and distribution of Pteronarcys californica Newport (Plecoptera) within streams. Great Basin Naturalist. 1973; 33: 218–220.

5. Surdick RF, Gaufin AR. Environmental requirements and pollution tolerance of Plecoptera. Environmental Protection Agency, Environmental Monitoring and Support Laboratory, Cincinnati, Ohio. 1978.

6. Kauwe JSK, Shiozawa DK, Evans RP. Phylogeographic and nested-clade analysis of the stonefly *Pteronarcys californicas* (Plecoptera: Pteronarcyidae) in the Western USA. Journal of the North American Bentholgical Society. 2004; 23: 824–838.

7. Myers LW, Kondratieff BC. Larvae of North American species of *Pteronarcys* (Plecoptera: Pteronarcyidae). Illiesia. 2017, 13:192-224. doi.org/10.25031/2017/13.16

8. Fore LS, Karr JR, Wisseman RW. Assessing invertebrate responses to human activities: Evaluating alternative approaches. Journal of the North American Benthological Society. 1996; 15: 212–231.

9. Freilich JE. Movement patterns and ecology of *Pteronarcys* nymphs (Plecoptera): observations of marked individuals in a Rocky Mountain stream. Freshwater Biology. 1991; 25: 379–394.

10. Needham JG, Claassen PW. A Monograph of the Plecoptera of North America North of Mexico. Entomological Society of America, Lafayette, Indiana. 1925.

11. Smith L. Studies of North American Plecoptera (Pteronarcinae and Perlodini). Transactions of the American Entomological Society. 1917; 43: 433–489. Available from: http://www.jstor.org/stable/25076980

12. Gaufin AR, Ricker ER, Miner M, Milam P, Hays RA. The stoneflies (Plecoptera) of Montana. Transactions of the American Entomological Society. 1972; 98: 1–161.

13. Brusven MA, Prather KV. Influence of stream sediment on distribution of macrobenthos. Journal of the Entomological Society of British Columbia. 1974; 71: 25–32.

14. Heinold BD. The mayflies (Ephemeroptera), stoneflies (Plecoptera), and caddisflies (Trichoptera) of the South Platte River Basin of Colorado, Nebraska, and Wyoming. M. Sc. Thesis, Colorado State University. 2010. doi: 10.13140/RG.2.2.24657.17761

15. DeWalt RE, Stewart KW. Life histories of stoneflies (Plecoptera) in the Rio Conejos of southern Colorado. Great Basin Naturalist. 1995; 55: 1–18.

16. Townsend GD, Pritchard G. Larval growth and development of the stonefly *Pteronarcys californica* (Insecta: Plecoptera) in the Crowsnest River, Alberta. Canadian Journal of Zoology. 1998; 76: 2274–2280.

17. Sheldon AL. Emergence patterns of large stoneflies (Plecoptera: Pteronarcys, Calineuria, Hesperoperla) in a Montana river. Great Basin Naturalist. 1999; 59: 169–174.

18. Fuller RL, Stewart KW. Stonefly (Plecoptera) food habits and prey preference in the Dolores River, Colorado. American Midland Naturalist. 1979; 101: 170–181.

19. Short RA, Maslin PE. Processing of leaf litter by a stream detritivore: Effect on nutrient availability to collectors. Ecology. 1977; 58: 935–938.

20. Erickson RC. Benthic field studies for the Windy Gap study reach, Colorado River, Colorado, fall 1980 to fall 1981. Prepared for The Northern Colorado Water Conservancy District, Municipal Sub-District. 1983.

21. Nehring RB. Stream fisheries investigations. Colorado Division of Wildlife, Federal Aid in Sportfish Restoration, Project F-51-R, Progress Report, Fort Collins. 1987.

22. Needham JG, Christenson RO. Economic insects in some streams of northern Utah. Logan, UT: Utah Agricultural Experiment Station Bulletin. 1927; 201: 1–36.

23. Baxter CV, Fausch KD, Saunders WC. Tangled webs: reciprocal flows of invertebrate prey link streams and riparian zones. Freshwater Biology. 2005; 50: 201–220. https://doi.org/10.1111/j.1365-2427.2004.01328.x

24. Muttkowski RA. The food of trout in Yellowstone National Park. Roosevelt Wild Life Bulletin. 1925; 2: 470–497.

25. Vinson M. A short history of *Pteronarcys californica* and *Pteronarcella badia* in the Logan River, Cache County, Utah. Utah State Bug Lab. Pteronarcyidae History Ongoing blog. Last updated 14 January 2008. Available from: https://www.usu.edu/buglab/Content/Files/salmonfly%20history.pdf

26. Birrell J, Nelson CR. Loss of the Giant Salmonfly *Pteronarcys californica* and changes in stonefly diversity in the Provo River, Utah (Plecoptera). Journal of Undergraduate Research, Brigham Young University. 2018. Available from: http://jur.byu.edu/?p=23170

27. Stagliano DM. Evaluation of salmonflies in Montana’s rivers: are statewide populations really declining? Montana Natural Heritage Program, Helena. 2010. 29 pp.

28. Nehring RB, Heinold BD, Pomeranz JF. Colorado River aquatic resources investigations. Colorado Division of Wildlife, Federal Aid Project F-237R-18, Final Report, Fort Collins. 2011. Available from: https://www.academia.edu/26701834/Colorado_River_Invertebrate_Investigations?auto=download

29. Ruse LP. Chironomid community structure deduced from larvae and pupal exuvia of a chalk stream. Hydrobiologia. 1995; 315: 135–142.

30. Foster SE, Soluk DA. Evaluating exuvia collection as a management tool for the federally endangered Hine’s emerald dragonfly, *Somatochlora hineana* Williamson (Odonata: Cordulidae). Biological Conservation. 2004; 118: 15–20.

31. Raebel EM, Merckx T, Riordan P, Macdonald DW, Thompson DJ. The dragonfly delusion: why it is essential to sample exuvia to avoid biased surveys. Journal of Insect Conservation. 2010; 14: 523–533.

32. DuBois RB. Detection probabilities and sampling rates for Anisoptera exuvia along river banks: influences of bank vegetation type, prior precipitation, and exuvia size. International Journal of Odonatology. 2015; 18: 205–215.

33. Richards DC, Rolston MG, Dunkel FV. A comparison of salmonfly density upstream and downstream of Ennis Reservoir. Intermountain Journal of Sciences. 2000; 6: 1–9.

34. Zippin C. The removal method of population estimation. Journal of Wildlife Management 1958; 22: 82–90.

35. Cooch E, White G. Program MARK: A Gentle Introduction. 17th edition. 2017. Available from: http://www.phidot.org/software/mark/docs/book/

36. Peterson JT, Thurow RF, Guzevich JW. An evaluation of multipass electrofishing for estimating the abundance of stream-dwelling salmonids. Transactions of the American Fisheries Society. 2004; 133: 462–475.

37. Riley SC, Fausch KD. Underestimation of trout population size by maximum likelihood removal estimates in small streams. North American Journal of Fisheries Management. 1992; 12: 768–776.

38. Branham JM, Hathaway RR. Sexual differences in the growth of Pteronarcys californica Newport and Pteronarcella badia (Hagen) (Plecoptera). Canadian Journal of Zoology. 1975; 53: 501–506.

39. Schultheis AS, Bootha JY, Vinson MR, Miller MP. Genetic evidence for cohort splitting in the merovoltine stonefly Pteronarcys californica (Newport) in Blacksmith Fork, Utah. Aquatic Insects. 2008; 187–195.

40. Huggins RM. On the statistical analysis of capture-recapture experiments. Biometrika. 1989; 76:133–140.

41. White GC, Burnham KP. Program MARK: survival estimation from populations of marked animals. Bird Study. 1999; 46:sup1: S120–S139.

42. R Development Core Team. R: A language and environment for statistical computing. 2012. R Foundation for Statistical Computing, Vienna, Austria. URL http://www.R-project.org/.

43. Saunders WC, Fausch KD, White GC. Accurate estimation of salmonid abundance in small streams using nighttime removal electrofishing: an evaluation using marked fish. North American Journal of Fisheries Management. 2011; 31: 403–415.

44. Vinson MR, Hawkins CP. Effects of sampling area and subsampling procedure on comparisons of taxa richness among streams. Journal of the North American Benthological Society. 1996; 15: 392–399.

